# The Prevalence of the Val66Met Polymorphism in Musicians: Possible Evidence for Compensatory Neuroplasticity from a pilot study

**DOI:** 10.1101/2020.12.23.424122

**Authors:** T.L. Henechowicz, J.L. Chen, L.G. Cohen, M.H. Thaut

## Abstract

It has been reported that MET carriers may express deficits in motor learning and neuroplasticity, possibly deterring musicianship. Here, we compared the prevalence of the Val66Met Brain-derived Neurotrophic Factor single nucleotide polymorphism (rs6265) in a sample of musicians (N=50) and an ethnically matched subset from the 1000 Human Genome Project (N-424). We report no differences in genotype or allele frequencies. Results are consistent with the view that hypothesized Met-dependent deficits in neuroplasticity are either mild or can be overcome by long-term practice.

## Introduction

Musicians serve as excellent models for studying neuroplasticity of the sensorimotor system, as music training uniquely involves long-term highly specific motor learning, often begins early in age, and involves error learning and multisensory feedback(1,2). However, genetic differences may influence both the likelihood of becoming a musician and the effects of music-induced plasticity(3). The Val66Met Brain-derived Neurotrophic Factor single nucleotide polymorphism (rs6265) (Val66Met BDNF SNP) is a common mutation present in 25-30% of the general population(4) that is associated with possible deficits in motor learning and neuroplasticity(5–7). Met-carriers show decreased activity-dependent secretion of pro-BDNF, which alters the secretion of mature BDNF, NMDA-receptor long-term potentiation (LTP), long-term depression (LTD), and the formation of inhibitory synapses(8). Due to the role of pro-BDNF in LTP processes, BDNF is a critical protein for learning, neuroplasticity, rehabilitation(9). The Val66Met BDNF SNP has been highlighted as an important gene for stroke rehabilitation due to Met-carriers’ poorer overall recovery and decreased response to brain stimulation interventions(9). Conversely, musicians have enhanced motor and sensory skills(10) and earlier onset of music training is associated with greater enhancements in sensorimotor learning(11). Long-term music training is a catalyst for neuroplasticity as musicians show numerous structural and functional brain changes (for reviews see 1,2). Therefore, the Val66Met polymorphism is a great candidate gene for investigating the relationships between music training and cortical plasticity. The objective of this study is to investigate the prevalence of the Val66Met BDNF single nucleotide polymorphism (SNP), a genetic mutation associated with deficits in neuroplasticity and motor learning, in a sample of musicians (N=50) compared to the general population (N=424) subset from the 1000 Human Genome Project). If musicians are less likely to have a genetic predisposition to deficits in motor learning and neuroplasticity plasticity deficits, the prevalence of the polymorphism (Val/Met genotype) in the musician sample would be significantly less than the sample from the general population

## Methods and Materials

Ethical approval was obtained from the University of Toronto Research Ethics Board; all participants provided written informed consent. For the control sample, genotype data were extracted from N=424 European samples from the 1000 Genomes Project. The 1000 Genomes Project has genotype data on 2318 individuals from 19 populations in 5 continental groups, generated on the Illumina Omni2.5 platform. We performed extensive quality control analyses and extracted a set of 1752 unrelated samples with high genotype quality. The subset included 119 CEU (Utah Residents with Northern and Western European Ancestry) samples, 110 TSI (Tuscan in Italy) samples, 95 GBR (British in England and Scotland) samples, and 100 IBS (Iberian in Spain) samples. We did not have demographic information for the HGP subset. For the control dataset, genotype data was used to infer sex for each individual.

We recruited a cohort of N=50 healthy musicians, currently enrolled in or recently completed a Bachelor’s degree in music performance (within 5 years) with four grandparents with descent from a European country (excluding Finland) matched to the HGP subset. We recruited European ancestries to control for the variation in the Val66Met prevalence between ethnicities(4). We recruited an equal number of males and females. For the musician sample, we recorded demographic variables of age, sex, degree program and year, primary instrument, years of primary instrument training, secondary instruments, and special musical achievements.

### Genotyping

Musicians provided a saliva sample using the DNA Genotek OG-500 kits. Genotyping for the SNP rs6265 (BDNF; Val66Met) was performed at The Centre for Applied Genomics, The Hospital for Sick Children, Toronto, Canada using a pre-designed TaqMan^®^ SNP Genotyping Assay (C__11592758_10, Life Technologies Inc., Carlsbad, CA, USA). The 10 ml reaction mix consisted of 5ml TaqMan Genotyping Master Mix (Life Technologies), 0.25 ml of 40X combined primer and probe mix, 2.75 ml water and 20-50 ng of DNA template. Cycling conditions for the reaction were 95°C for 10 min, followed by 40 cycles of 94°C for 15 sec and 60°C for 1 min. Samples were analyzed using the ViiA™ 7 Real-Time PCR System and analyzed using ViiA™7 software.

### Statistical Analysis

Statistical analyses were conducted in R. Due to the small cell count (<5), Fisher’s exact test was used to assess significant differences in genotype frequencies and the Chi-square test was used to detect significant differences in allele frequencies (two-tailed and alpha = 0.05). Due to the small count for the Met/Met genotype (N=1), we compared the demographic variables between Val/Val and Met-carriers (Val/Met and Met/Met). We conducted two-sample t-tests for age, total years of training, and years of training on the primary instrument. We did not stratify by instrument type due to sample size.

## Results

The mean age of the musician sample was 21.8± 3.5 with 11.7± 4.7 years of training on their primary instrument and 14.3± 3.6 years of total music training (See Table.1). The musicians included instrumentalists, woodwind (N=16), brass (N=7), strings (N=15), and percussion (N=4) as well as keyboard players (N=8). Voice majors were not because singing may involve different neural processes from instrumental music training, where motor learning involving the upper limbs are critical to performance. N=37 musicians had pre-university awards (competitions, scholarships, festivals), N=13 musicians received awards while in university, and N=22 musicians had professional ensemble placements. The HGP subset containd N=210 Males, N=207 Females, and N=7 undetermined samples. Since the HGP subset was a sample of the general population, the HGP subset may include some musicians but we assume that this would be a small percentage. The musician sample and the HGP subset were in Hardy-Weinberg equilibrium (p=0.24 for musicians, p=0.75 for HGP).

**Table 1.** Demographic Variables for the Musician Sample by Carrier Status

| *Variable* | *Val/Val (N=29)* | *Met-Carriers (N=21)* | *Total (N=50)* |
| --- | --- | --- | --- |
| Age (at time of data collection) | 21.8 ± 3.0 | 21.7 ± 4.2 | 21.8 ± 3.5 |
| Age of start for musical training | 8.1 ± 3.1 | 6.5 ± 3.9 | 7.4 ± 3.7 |
| Sex (M/F) | 15/14 | 8/13 | 25/25 |
| Handedness (R/L) | 29/0 | 18/3 | 47/3 |
| Total Years of Training | 13.7 ± 3.0 | 15.3 ± 4.1 | 14.3 ± 3.6 |
| Years of Training on Primary Instrument | 10.9 ± 3.9 | 12.0 ± 5.4 | 11.7 ± 4.7 |
| Number of musical achievements | 3.1 ± 1.1 | 3 ± 1.3 | 3.06 ± 1.2 |

The results revealed that there were no significant differences in genotype frequencies (p=0.6447) and allele frequencies (p=0.8513) (Figs 1 and 2). There were no significant differences between Val/Val and Met-carriers for age (t = 0.074043, df = 34.076, p-value = 0.9414), total years of music training (t = −1.5248, df = 34.546, p-value = 0.1364), years of training on primary instrument (t = −1.4926, df = 34.623, p-value = 0.1446), and age of start of music training (t = −1.5945, df = 39.881, p-value = 0.1187). The number of early starters (before 6.5 years old) versus late starters (after 6.5 years old) were not significantly different (X-squared = 3.6872, df = 1, p-value = 0.05483).

**Figure 1.**
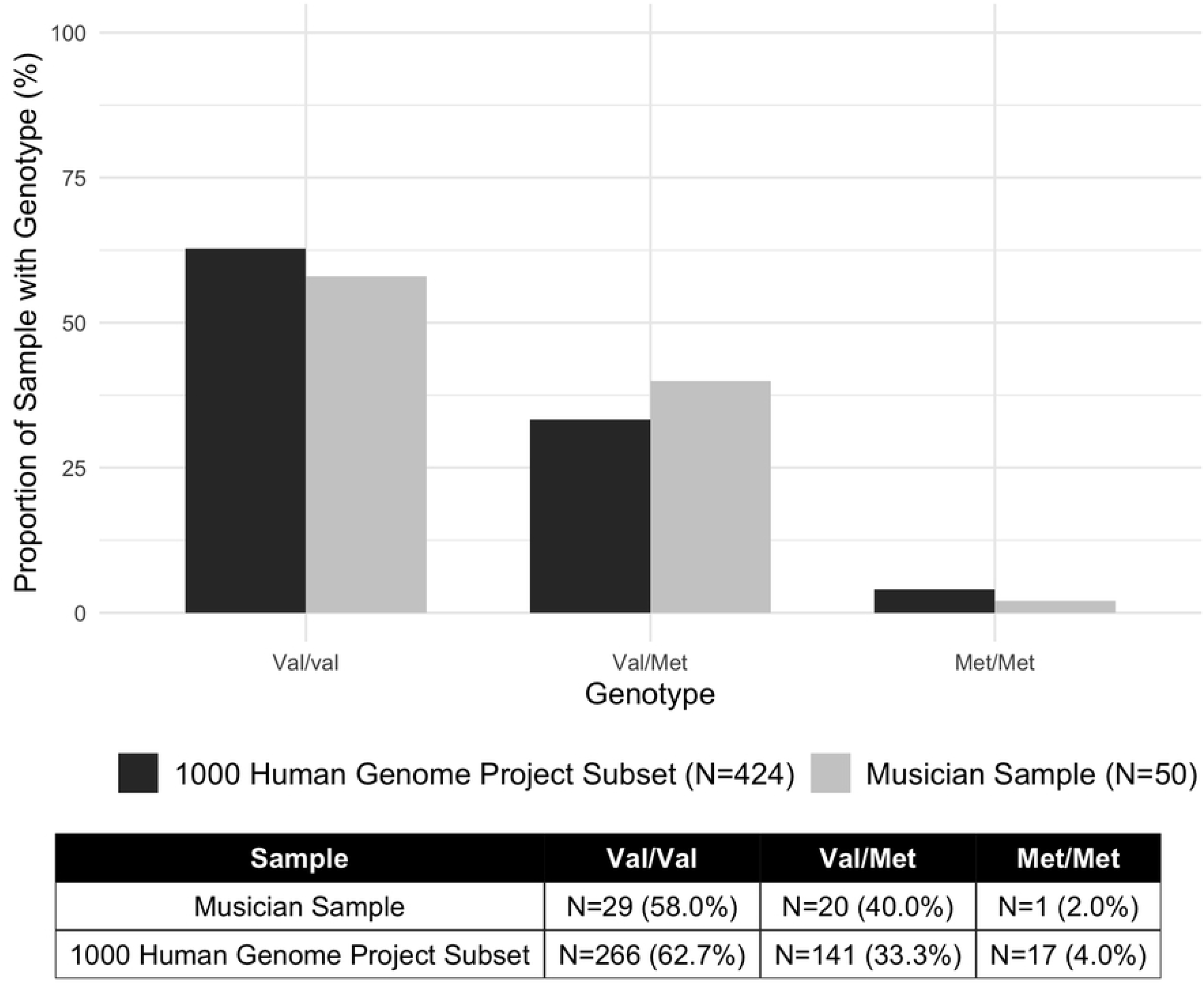
Genotype Frequency Distributions in the Musician Sample and 1000 Human Genome Project Subset.

**Figure 2.**
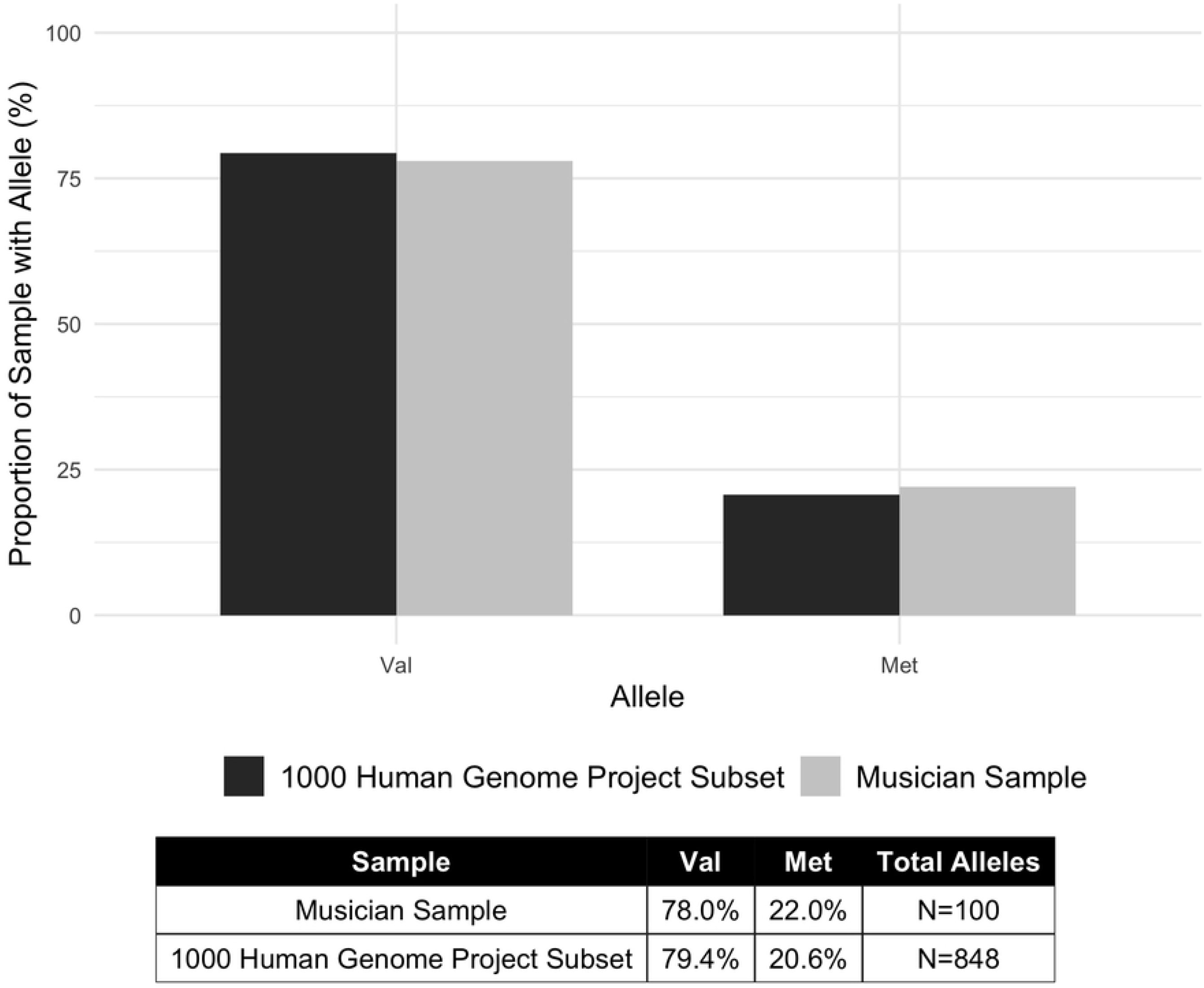
Allele Frequency Distributions in the Musician Sample and 1000 Human Genome Project Subset.

## Discussion

Long-term and intensive music training induces structural and functional brain changes, and enhances short-term plasticity(1,2,12,13). The presence of the Val66Met BDNF polymorphism (Val/Met or Met/Met) genotype is associated with altered cortical plasticity(5) and deficits in motor learning(6). The objective of this study was to assess the prevalence of the Val66Met polymorphism in musicians compared to the general population. We report that they were similar. The Val66Met polymorphism does not overtly limit musicianship. It is possible that intense training beginning early in life and involving long-term deliberate practice(1) required for successful musicianship overcomes inherent MET-dependent deficits in the response to motor training(14). This view is consistent with the finding that practice improves Met-dependent deficits in non-musicians(15). A meta-analysis of N=55 studies revealed that single sessions of acute aerobic exercise increase activity-dependent BDNF which may have enhancing effects for motor learning(16). Thus, music training, an intense motor activity, may also increase activity-dependent BDNF. Differences in brain connectivity between musicians and controls that correlate with years of practice(17) could represent one neural substrate supporting this possibility. Alternatively, it is possible that Met-dependent deficits alone are mild and not enough to elicit training-dependent deficits in musicians, requiring for example other genetic abnormalities to express(3). Although the genetics of musical motor timing have been explored(18), the genetics of musical motor learning are not known. Future studies, involving larger “n”s and possibly other plasticity probes could inform on the impact of the Met-Met anomaly, present in only one musician in our sample, on musicianship.

In neurorehabilitation, the Val66Met polymorphism disrupts motor plasticity in stroke patients and may hinder motor function recovery(9). However, Neurologic Music Therapy [NMT] interventions for stroke patients induce cortical changes in the organization of the sensorimotor cortex and improve motor function(19,20). Future NMT clinical trials should consider the predictor of Val66Met polymorphism status, which may bring insight into the role of BDNF-dependent plasticity in music-based interventions.

## Acknowledgements

Thank you to the team at the Centre for Applied Genetics, Sick Kids Hospital, Toronto for genotyping and statistics consultations.

## Author Contributions

M.T and L.C. conceived of the study rationale and design. T.H. methodologies, carried out data collection, conducted statistical analysis, reviewed literature, and wrote the first draft of the manuscript. M.T. supervised all data collection. J.C. provided feedback at numerous stages of the project. All authors provided feedback and contributed to manuscript writing and revisions.

